# Information processing dynamics in neural networks of macaque cerebral cortex reflect cognitive state and behavior

**DOI:** 10.1101/2021.09.05.458983

**Authors:** Thomas F. Varley, Olaf Sporns, Stefan Schaffelhofer, Hansjörg Scherberger, Benjamin Dann

## Abstract

One of the essential functions biological neural networks is the processing of information. This comprises processing sensory information to perceive the environment, up to processing motor information to interact with the environment. Due to methodological concerns, it has been historically unclear how information processing changes during different cognitive or behavioral states, and to what extent information is processed within or between the network of neurons in different brain areas. In this study, we leverage recent advances in the calculation of information dynamics to explore neural-level processing within and between the fronto-parietal areas AIP, F5 and M1 during a delayed grasping task performed by three macaque monkeys. While information processing was high within all areas during all cognitive and behavioral states of the task, inter-areal processing varied widely: during visuo-motor transformation, AIP and F5 formed a reciprocally connected processing unit, while no processing was present between areas during the memory period. Movement execution was processed globally across all areas with a predominance of processing in the feedback direction. Additionally, the fine-scale network structure re-configured at the neuron-level in response to different grasping conditions, despite of no differences in the overall amount of information present. These results suggest that areas dynamically form higher-order processing units according to the cognitive or behavioral demand, and that the information processing network is hierarchically organized at the neuron-level, with the coarse network structure determining the behavioral state and finer changes reflecting different conditions.

**Significance Statement:** What does it mean to say that the brain “processes information?” Scientists often discuss the brain in terms of information processing – animals take in information from their environment through their senses, and use it to make decisions about how to act in the world. In this work, we use a mathematical framework called information theory to explore how signals from the environment influence brain activity, and how brain activity in turn informs on behaviors. We found that different brain regions processed information in dynamic and flexible ways, with signals flowing up and down the hierarchy of sensory-motor depending on the demands of the moment. This shows how “computation” in the brain can reflect complex behaviors and cognitive states.

## 1 Introduction

Animal nervous systems are often described as “information processing engines:” organisms take in information about the world around them through sensory organs, learn statistical regularities in the incoming information, and use those to navigate and interact with their environment. As a distributed network of neurons, information processing takes place both within and between many different specialized brain regions. These processes can be categorized as feedforward processing (from primary sensory areas to motor areas) and feedback processing (from motor areas back to sensory areas, see Fig. 1). Despite a growing number of studies demonstrating the involvement of multiple areas in different behavioral processes [1, 2, 3, 4], it remains unclear to what degree information processing in the brain’s network of neurons takes place restricted to within specific areas, as opposed to multi-areal feedforward and feedback interactions. Furthermore, it is unclear whether the intra- and inter-areal processing network structure at the neuron-level is static or changes dynamically in response to the demands of particular cognitive or behavioral states (Figure 1B).

**Figure 1:**
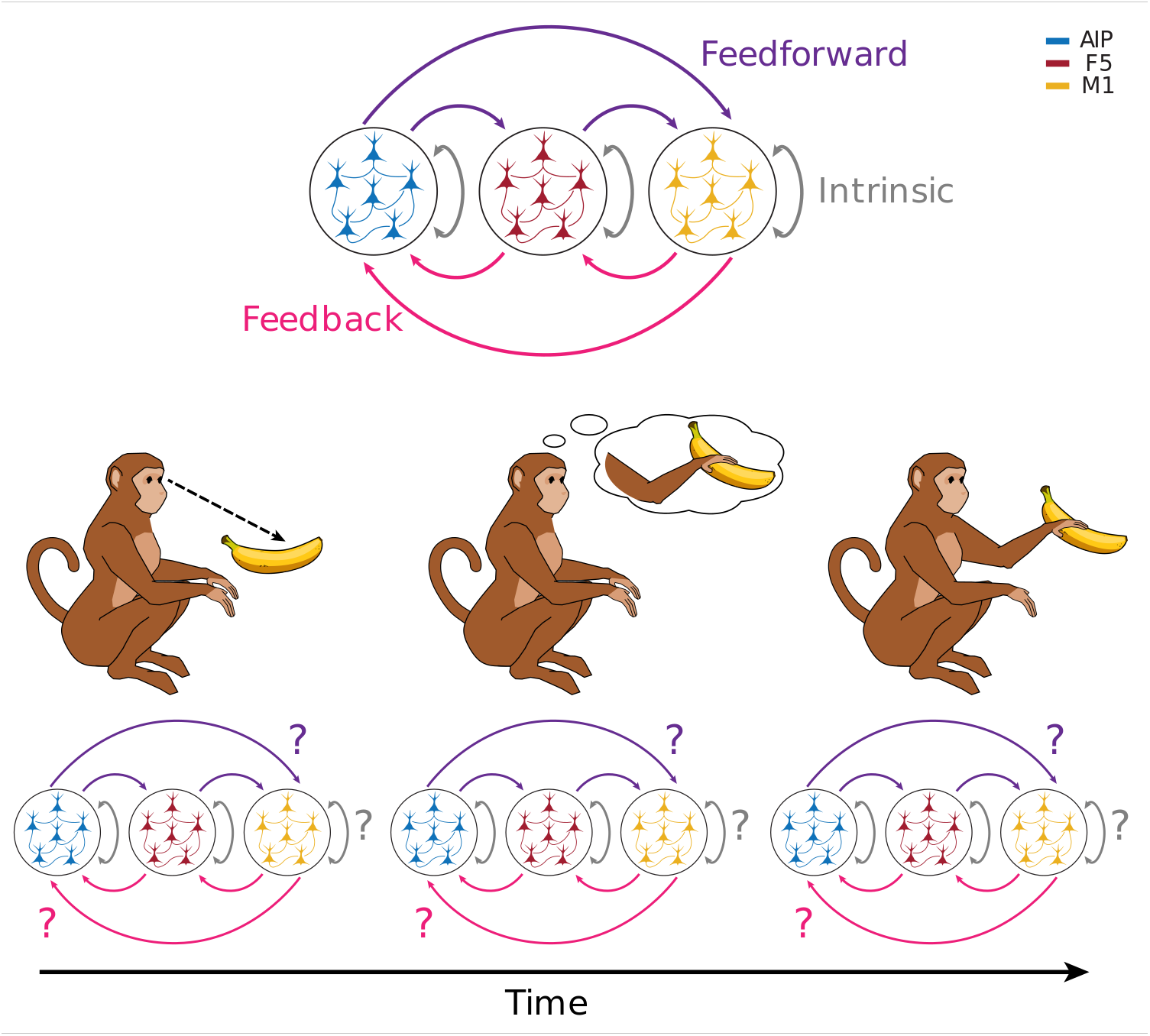
Feedforward and feedback computation in the brain. Information processing associated with object recognition and motor planning takes place in multiple, distinct brain areas of the sensory-motor network. How information processing is distributed over those areas, and how that distribution “updates” as the task changes is unknown. Here we delineate three cortical areas (M1, AIP, and F5), and the various kinds of intrinsic, feedforward, and feedback flows of computation and information possible between them. Over the course of the experiment, the degrees to which these different modes dominate the overall computational structure may change.

Given the technical limitations of whole-brain recording, it is useful to focus on subsets of areas that display a range of intrinsic, feedforward, and feedback modes of processing to fully understand how they relate. A natural system is the set of brain areas known to be involved with perception- and movement execution. In macaque monkeys, the fronto-parietal grasping network specifically has been shown to be strongly involved in visuo-motor transformations and the execution of grasping movements and is composed of the anterior intraparietal area (AIP), the ventral premotor cortex (F5), and the primary motor cortex (M1) [5, 6, 7, 8]. Moreover, several studies demonstrated that AIP and F5 are strongly reciprocally connected, as are F5 and M1 [9, 10, 11, 12, 13] suggesting this network to be a compelling candidate to study intra- and inter-areal processing.

When studying activity of simultaneously recorded neurons, however, information processing is often vaguely defined. The mathematical fields of information theory and the theory of information dynamics offers a general framework to study information flow and processing. Information dynamics breaks the concept of information processing into three parts[14]:

**Information storage**: The degree to which the past activity of a neuron informs on it’s future, (e.g. LTP or LTD.) [15],
**Information transfer**: The degree to which the past of a source neuron informs a target neuron’s future, (e.g. synaptic communication) [16, 17],
**Information modification**: The “non-linear” computation that occurs when a neuron integrates distinct streams of information into something greater than the sum of its parts. [18, 19, 20, 21].

These three dynamics can be formalized using information theory [22] (see Sec. 1.1). Previous work using information dynamics to study recorded neuronal networks found that the degree of information modification changes during development [21], and in the same developmental windows particular patterns of information transfer are “locked-in” [23]. Furthermore, the capacity for information modification is heterogeneously distributed across neurons of the network; concentrated in high-degree, rich-club neurons [24, 25, 26]. Information transfer [27] has been applied to a variety of neural recordings (see [16] for a comprehensive review) and allows researchers to estimate effective network models of interacting neurons. Finally, active information storage has provided insights into stimulus response and preferences in visual processing systems [15].

Despite the wealth of analyses that have been done using information theory to investigate neural activity, information theory has not been less frequently applied to the problem of behavior-related information processing at the neural network-level. Similarly, differences in neural information dynamics have not been thoroughly investigated within and between areas during complex behaviors. Much of the above-cited work has been done in neural cultures or anesthetized animals, as opposed to behaving organisms interacting with a complex environment. To address this gap, we examined the information dynamics and the associated effective network structures of neural populations from three cortical areas (AIP, F5 and M1) in the fronto-parietal grasping network of three macaques. During recordings, monkeys performed a delayed sensory-motor transformation task involving the transformation of visual information into movement plans, the memorization of these movement plans, and finally the required processing to execute one of two grasping movements (for details, see [28]).

With these data, we can estimate the neuron-level information dynamics in different cognitive and behavioral states, allowing us to directly assess the relationship between information dynamics and complex behaviors. Furthermore, by casting patterns of information transfer as effective connectivity networks, we can examine the degree of intrinsic, feedforward and feedback information processing within and between the three areas, and how changes in behavior alter the information dynamics within and between them. We found that different behavioral states were associated with significant re-configurations of the global effective network structure, with movement in particular being associated with an increase in the overall information flowing through the system, and an increase in the amount of synergistic information processing. Within and between the different areas, we found that: (1) information processing was strong both within and between AIP and F5 during visuo-motor transformation, (2) during memorization of movement plans, information processing remained high within areas but strongly decreased between AIP and F5, and (3) globally, information processing was strongest during movement execution in decreasing order from M1 to F5 to AIP both within and between areas, showing a more pronounced pattern in the feedback direction. Moreover, in all three behavioral states, the network-wide patterns of information storage and transmission were different for each of the two different grasping conditions. Together, these findings suggest that the information processing structure at the neuron-level changes depending on the behavioral state and task condition, whereas state changes are predominantly associated with changes in information processing between areas, condition changes are predominately associated with fine-scale reconfiguration of the network structure.

### 1.1 Basic Theory of Information Dynamics

A natural mathematical framework for assessing these computational dynamics is *information theory* [22, 14], which describes how the activity of interacting elements of a complex system constrain the space of possible states of the whole system. A significant advantage of information theory is that, being based only on marginal, joint, and conditional probabilities of events, it is both model-free and sensitive to non-linear relationships between interacting elements [29]. Information theoretic analysis is then “epistemically modest”, as it does not require presupposing particular generating functions or relationships. This makes it ideal for complex, nonlinear systems, such as networks of neurons of brains, where the underlying generative dynamic is unknown and nonlinearities can play a key role [30].

Information theory quantifies the how knowledge of a variable, or set of interacting variables, reduces the uncertainty of an observer watching the system. The central object of study is the *entropy* of a random variable, which quantifies an observer’s uncertainty about the state of a variable under study. For a discrete random variable *X*, with support set 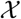, the entropy *H*(*X*) is canonically given as:

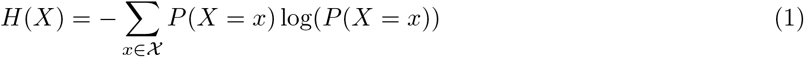

When measured in bits (I,e. The base of the logarithm is two), the entropy of *X* quantifies the minimum number of yes/no questions required to completely specify the state of *X* with total certainty. Given two variables *X* and *Y*, we can calculate the *mutual information* between the two as the degree to which knowing the state of one variable reduces our uncertainty about the state of the other. Formally:

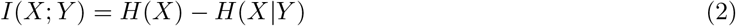

Where *H*(*X*|*Y*) is the conditional entropy of *X* given *Y*. The difference between our initial uncertainty about *X* and the remaining uncertainty after accounting for *Y* is the amount of uncertainty about *X* that *Y* resolves.

For temporally extended processes, we can break down the total information structure of the system in three information dynamics: active information storage, information transfer, and information modification. Information storage quantifies the degree to which knowing the past of a variable, or set of variables, decreases our uncertainty about its immediate future:

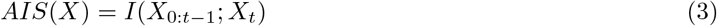

Where *X*_0:*t*−1_ is the joint state of all past states of variable *X* and *X_t_* is the immediate next state. In the context of neurons, a process that generates non-trivial AIS might be hyperpolarizing a neuron, which will reduce the probability that it will fire in the immediate future [15].

For sets of interacting variables, we can generalize the active information storage to quantify how much information is “transferred” from a source neuron to a target neuron by determining how much knowing the past of a prospective “source” variable *X* reduces our uncertainty about the future of a “target” variable *Y* (above and beyond the information stored in the target):

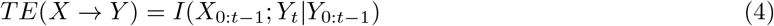

*TE*(*X* → *Y*) is referred to as the transfer entropy from *X* to *Y*. The canonical example here is that of synaptic communication: knowing that an excitatory pre-synaptic neuron spiked increases our certainty that the post-synaptic neuron will spike in the near future. Importantly, the I/O functions of neurons themselves are highly complex [31], and so the future behavior of the neuron may depend on the *collective* behaviour of all the upstream sources, which can be quantified with the multivariate transfer entropy:

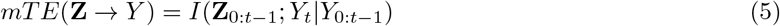

where **Z** is an ensemble of neurons. To recover the effect of a single source on the target *in the context of all other informative sources*, we use the conditional multivariate transfer entropy [32, 33, 34]:

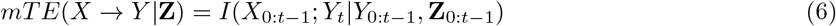

Finally, the third information dynamic, information modification, refers to information produced when multiple incoming “streams” intersect and are non-trivially changed. This is operationalized as the “synergy” [18, 35] (for mathematical details see Sec. 4.9). Briefly, synergy can be understood as the information provided by a set of sources that cannot be extracted from any simpler combination (i.e. the marginals). While the existence of synergy in biological neural networks is extremely well documented [21, 24, 25, 26, 36], its exact biological or behavioral significance remains unclear.

## 2 Results

### 2.1 Behavioral task and single neuron recordings

To study behavior-related neural information processing within and between areas, we utilize data recorded from three monkeys (S, Z, and M). Monkeys were trained to perform a delayed grasping task (Fig. 2 A). The task was divided into four epochs: (1) an initial fixation epoch, which was the same for all conditions, (2) a cue epoch, in which the monkeys were either instructed or free to choose to grasp a target with one of two possible grip types (power or precision grip; monkey M was only trained to performed the instructed context), (3) a memory epoch, in which the monkeys had to prepare and remember the corresponding grip-type, and (4) a movement epoch, in which the monkeys performed the corresponding grip-type (for details, see Materials & Methods). While monkeys performed the task, we simultaneously recorded large populations of well isolated neurons from the ventral premotor cortex (area F5), the anterior intraparietal area (AIP), and for one monkey from the hand area of the primary motor cortex (M1) (Fig. 2 B). For this purpose, two 32-channel microelectrode arrays were chronically implanted per area resulting in 128 recorded channels for monkey S and Z and 192 recording channels for monkey M. For all following analyses, we used 4 recording sessions from monkey S with on average 82.8 +/- 5.7SD neurons, 3 sessions from monkey Z with on average 53 +/- 5.6SD neurons, and 3 sessions from monkey M with on average 134.3 +/- 23.7SD neurons.

**Figure 2:**
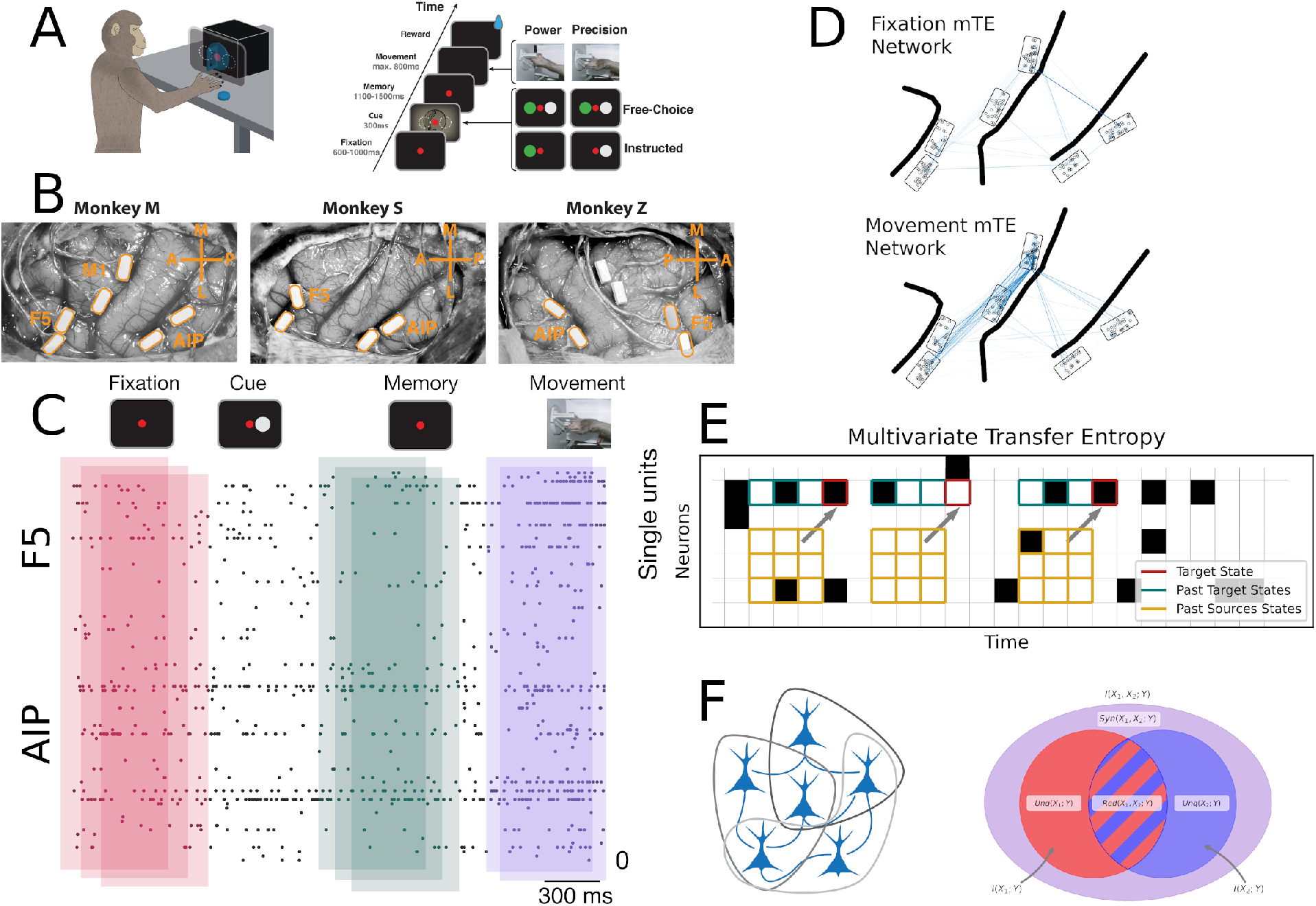
Electrode array implantation and behavioral task structure. **A** The macaque was placed in front of a screen on which distinct symbols can be displayed. The macaque had been trained to, depending on the symbol that appeared, execute one of two distinct grip types (precision grip or power grip). The structure of the task was that each trial would begin with an epoch of passively fixating a red disc on a screen, followed by the onset of the cue, another epoch of the fixation cue, where the appropriate movement was prepared and memorized, and finally the signal to execute the appropriate grip type. **B** Pictures of implanted floating micro-electrode arrays in monkey M (left), monkey Z (middle) and monkey S (right). Monkeys were implanted with 2 floating Microprobe arrays per area (4-6 total), in the areas AIP and F5 and M1 of the fronto-parietal grasping network. **C** The sliding-windows approach: over the course of the entire sequence, a window of 800 ms was incremented by 100 ms steps, and for each window, the information theoretic analyses were repeatedly performed. **D** The structure of two representative multivariate transfer entropy networks for two different epochs (fixation and movement), projected onto the cortical arrays. **E** A cartoon explaining the intuition behind the multivariate transfer entropy. We should note that the actual transfer entropy algorithm takes a continuous “sliding windows” approach (not to be confused with the sliding windows reported in panel C) - in this cartoon, only a subset of “windows” are visualized for the purposes of building intuition. **F** A visualization of the triadic partial information decomposition: for every set of three neurons arranged into two sources informing on a singlet target, we can decompose the joint mutual information from both sources into redundant, unique, and synergistic components.

### 2.2 Information Dynamics

Behavior dependent changes in information dynamics were estimated by computing AIS, conditional mTE and triadic synergy between all simultaneously recorded neurons (see Materials & Methods). All three measures were computed over the time course of the task with a sliding window of 800ms and separatef by grasping condition (incremental step size of 100ms). The result is an array of windows: 23 time windows x 2 grasping conditions; (Fig. 2 C). Note that in order to reduce the influence of behavior dependent firing rate changes, the AIS and the mTE measures were significance tested with conservatively estimated surrogate data (shuffling spikes within 25ms windows and recomputing the corresponding measure; for details, see Materials & Methods). In addition, all values (AIS, mTE, and synergy) were normalized by dividing by the Shannon entropy of the receiver neuron, following [24, 25, 26], which provides a control for variable firing rates.

For a first assessment of differences in information dynamics during behavioral changes, we compared the average values of the three basic normalized information dynamics per behavioral epoch across all neurons and areas (Figure 3 A-C). Kruskal-Wallis analysis of variance found a small, but significant difference between all four behavioral conditions for AIS (*H* = 67,*p* = 4.27 × 10^−15^). The effect size change in AIS values, however, was very small: the highest value was 0.014±0.024 (in the fixation condition), while the smallest was 0.012 ±0.021 (Cohen’s D = 0.1). This suggests that during the movement condition, the degree to which information is being “stored” in individual neurons is decreased, possibly in favor of information flowing from sources to targets. There was a stronger, significant difference in mTE between behavioral states (*H* = 1555.649, *p* < 10^−20^). Here the lowest average mTE was observed in the fixation period (0.005 ± 0.007) and the highest was observed in the movement period (0.013 ± 0.02, Cohen’s D = 0.46). The strongest significant difference was found with synergy (*H* = 1520.213, *p* < 10^−20^), with the smallest synergy being seen in the memory condition (0.001 ± 0.002) and the highest, as expected, being seen in the movement state (0.002 ± 0.003, Cohen’s D = 0.54). Time-resolving information dynamics over all time windows and across conditions (Fig. 3, D-F) show that the distributions of all three information dynamics were largely similar within behavioral states and between conditions. The only exception was the memory state, where the mTE and the synergy started to increase before the onset of movement. This is unsurprising since movement-related processing in cortical motor areas precedes movement onset by at least the amount of neuronal distance to the arm and hand [37, 38].

**Figure 3:**
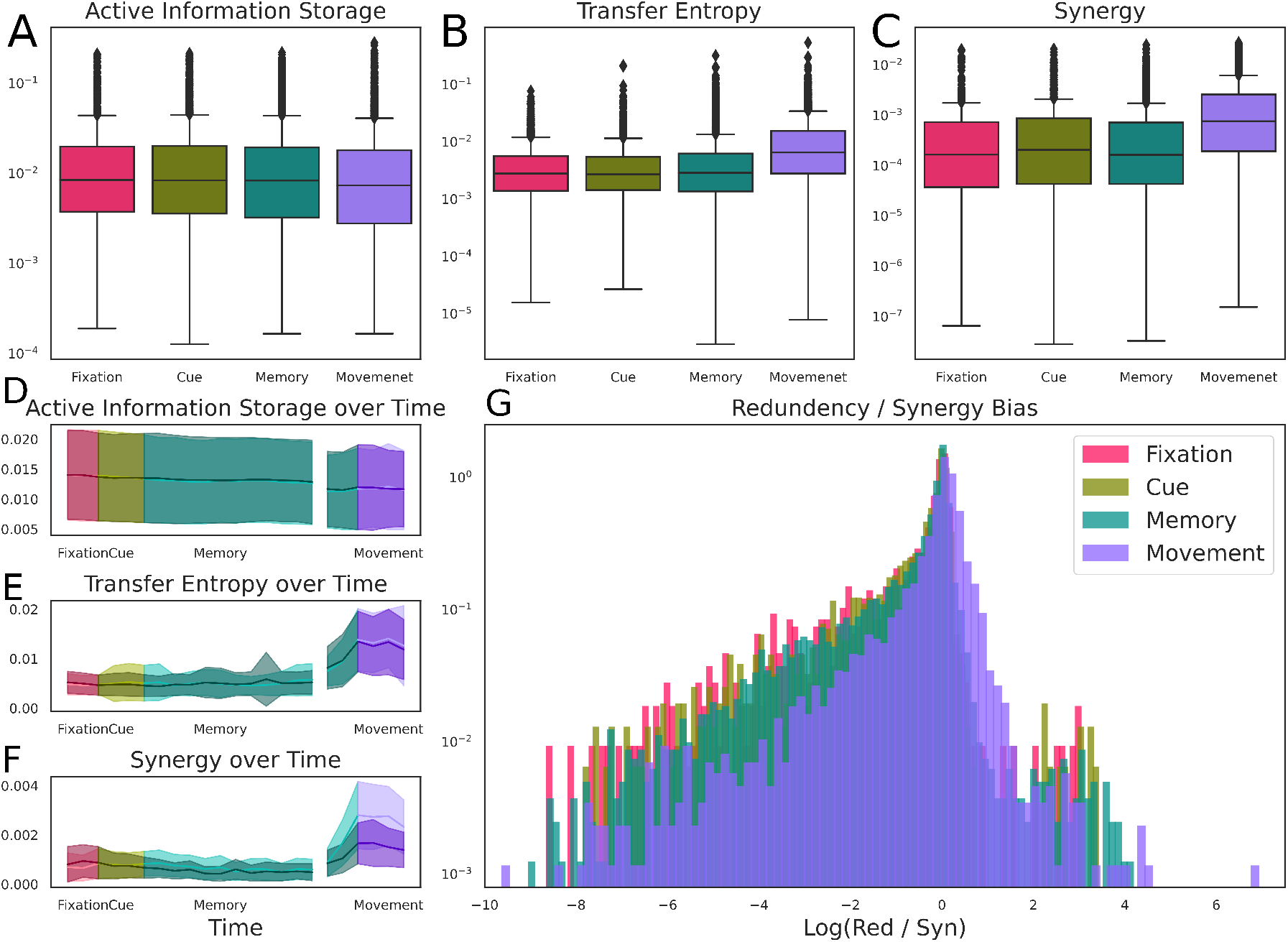
Information dynamics & behavioral states: **A - C**. The differences in AIS (left), mTE (middle), and synergy (right) between the four behavioral states. There is no significant difference for AIS over time, however mTE and synergy show dramatic increases during the movement epoch. **D - F** Sliding windowed, time-resolved differences between conditions. These figures provide a more finely time-resolved picture of how information dynamics vary between behavioral states and grasp conditions. For example, we can see that the mTE and synergy were starting to increase prior to the visible onset of movement. The shaded region accounts of ±one standard error and the two curves (dark and light) correspond to Grip Type A and B respectively. **G** The synergy / redundancy bias between behavioral states plotted as histograms. All the states are, on average, synergy-biased, although the onset of the movement behavior is associated with a distinct increase in the proportion of information that is redundant.

#### 2.2.1 Redundancy / Synergy Bias

The measure of information modification (synergy) also reveals an additional set of information processing “modes”, formalized by the partial information decomposition (PID) framework [39, 40], which reveals two different ways information can flow through a system: a redundancy-dominated mode, where information is duplicated and sent through many channels simultaneously, or a synergy-dominated mode, where information is distributed over higher-order combinations of multiple channels. These distinct modes may be thought of as different “ways” that the nervous system can distribute, process, and represent information.

We can assess which of these modes dominates the overall joint mutual information *I*(*S*_1_, *S*_2_; *T*) by normalizing the value of each atom (i.e. the portion of the joint-MI that is redundant, unique, or synergistic) by the total mutual information. Note that this normalization is different from the one reported above (where the value of every information dynamic is normalized by the entropy of the receiver neuron). We found that, in all behavioral states, the information flow was synergy dominated (see Fig. 3G), and that for the first three behavioral states (fixation, cue, memory), the values of synergy, redundancy the ratio of the two remained relatively constant. The transition from memory to movement, however, was associated with a marked increase in the relative redundancy of information flow (Δ = −74.4 ± −27.3%, U=10084455.0, p< 10^−10^). No other transition (fixation →cue, or cue →memory) was significant, despite the very large number of samples in each distribution. This indicates that the transition from “cognitive” states to motor states is associated with a particular increase in the relative dominance of redundant information communicated within the network.

### 2.3 Inter-Areal Analysis

Given the behavior-dependent changes in information dynamics over the course of the task, we next examined the degree to which the different types of information processing change within and between areas. To compare information dynamics of the different types within and between areas, we computed the average mTE and synergies for all neuron pairs *within* each area and also *between* all pairs of areas separately in the forward and backward directions. Depicted in Figure 4 are the average mTE and synergy dynamics for all areas and area combinations across all neuron pairs, recording sessions and monkeys. The dynamics of both synergy and mTE are highly similar, consequently we will not describe them separately in this section and instead refer to the overall pattern of “information processing.” Overall information processing was highest within areas, followed by directly connected areas and weakest between AIP and M1. Encouragingly, the overall information processing reflects the known anatomical connection strength, which we interpret as providing validation of our methods.

**Figure 4:**
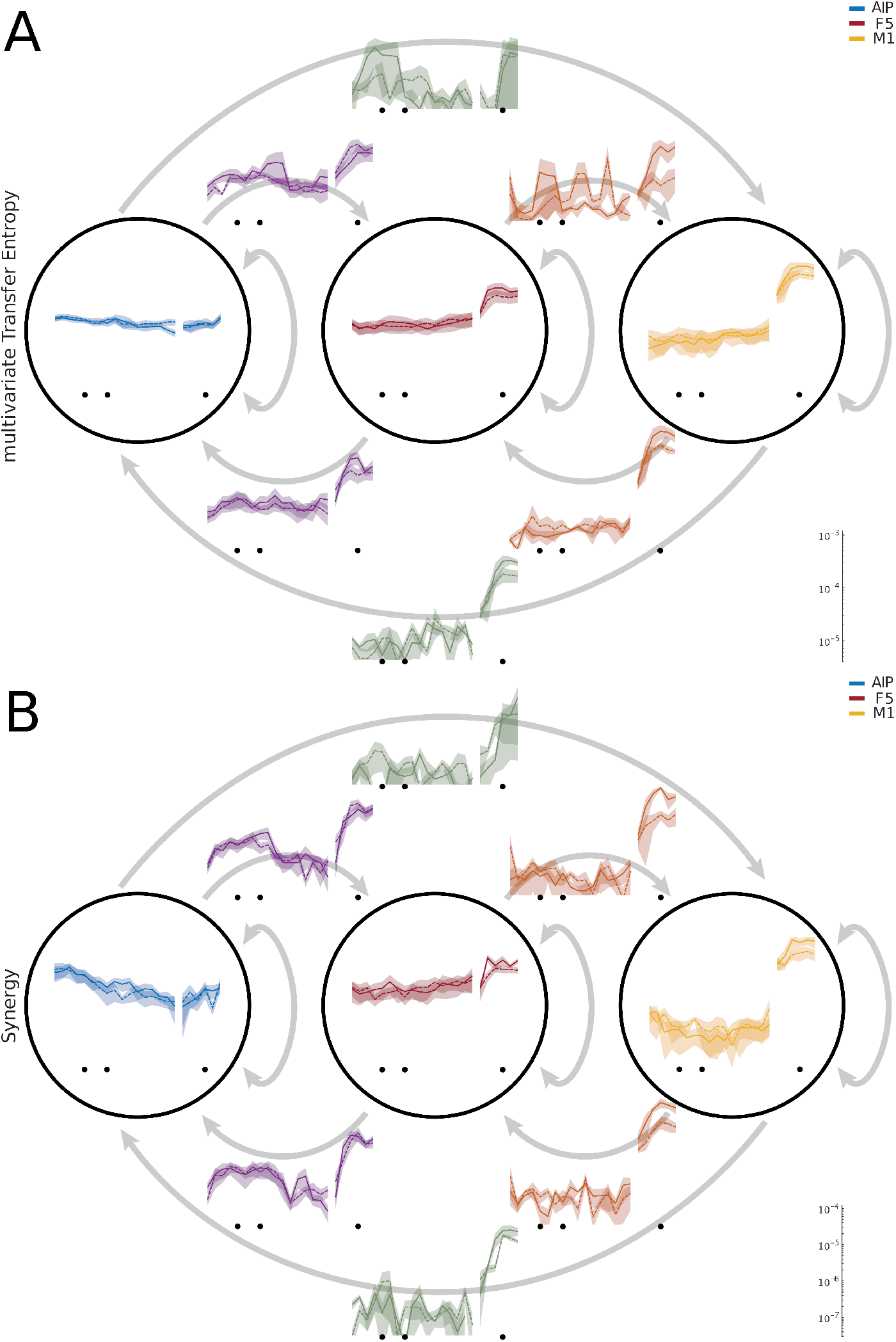
The information dynamics of mTE and synergy within and between the areas AIP, F5 and M1. Since both mTE and synergy are directed measures, the information dynamics between areas can be separated into the feedforward and feedback directions. **A** The strength of the normalized multivariate transfer entropy within areas (indicated by the circles) and between areas (where the source and the target are two different area. The arrow indicates the feedforward and feedback direction from AIP to F5 to M1). We can see that the degree of information processing varies greatly between areas for different behavioral states. **B** The same plot as A, but for normalized triadic synergy.

In AIP, despite an overall high level of information processing, there was surprisingly little change over the course of the task. In F5, in contrast, information processing increased during movement execution after remaining largely constant during the more “cognitive” behaviors. The increase in information processing during movement was even higher in M1, while information processing was lower during all other epochs in comparison to the other two areas.. Between areas, the amount of information processing was surprisingly similar in both the feedforward and feedback directions, suggesting processing in both directions to be of equal importance for the visuo-motor transformations and the execution of grasping movements. There were, however, significant differences between the different inter-area combinations. Area AIP and F5 showed increased inter-areal information processing around the cue epoch, in contrast to processing within both areas (which remained largely constant), suggesting that the processing between these areas is of particular importance for the transformation of visual information into movement plans. Apart from the already-high level of information processing within areas, no inter-area connection showed an increased level of information processing during the memory epoch, suggesting that the act of “holding information in memory” is an area specific, rather than a global process. In contrast, all inter-area combinations showed a strong increase in information processing during movement execution, suggesting the control of movement execution to be a rather globalized process not limited to M1.

In addition to the condition-dependent changes in synergistic processing, the PID framework allows us to also explore the joint mutual information, the redundant information, and the unique information atoms. For a better comparison between the different types of information processing, we averaged information processing for the three most important behavioral epochs of interest: cue, memory and movement to assess how different types of information flowed through the system. Figure 5 shows the average amount each information atom for all areas and inter-area pairs. As we have already observed in the comparison of mTE and synergy, temporal dynamics of all information types are generally similar. On closer inspection, however, small differences between the information types become apparent. During the visuo-motor transformation around the cue epoch, synergistic and redundant information processing between AIP and F5 were stronger relative to the three other types of information processing. During memory, low levels of mTE, synergistic, and redundant information processing were present between AIP and F5.. Presented in this way, the aforementioned global information processing patterns during movement execution become clearly apparent. Furthermore, it becomes clear that all types of information processing during movement execution are more intense in the feedback direction then in the feedforward direction.

**Figure 5:**
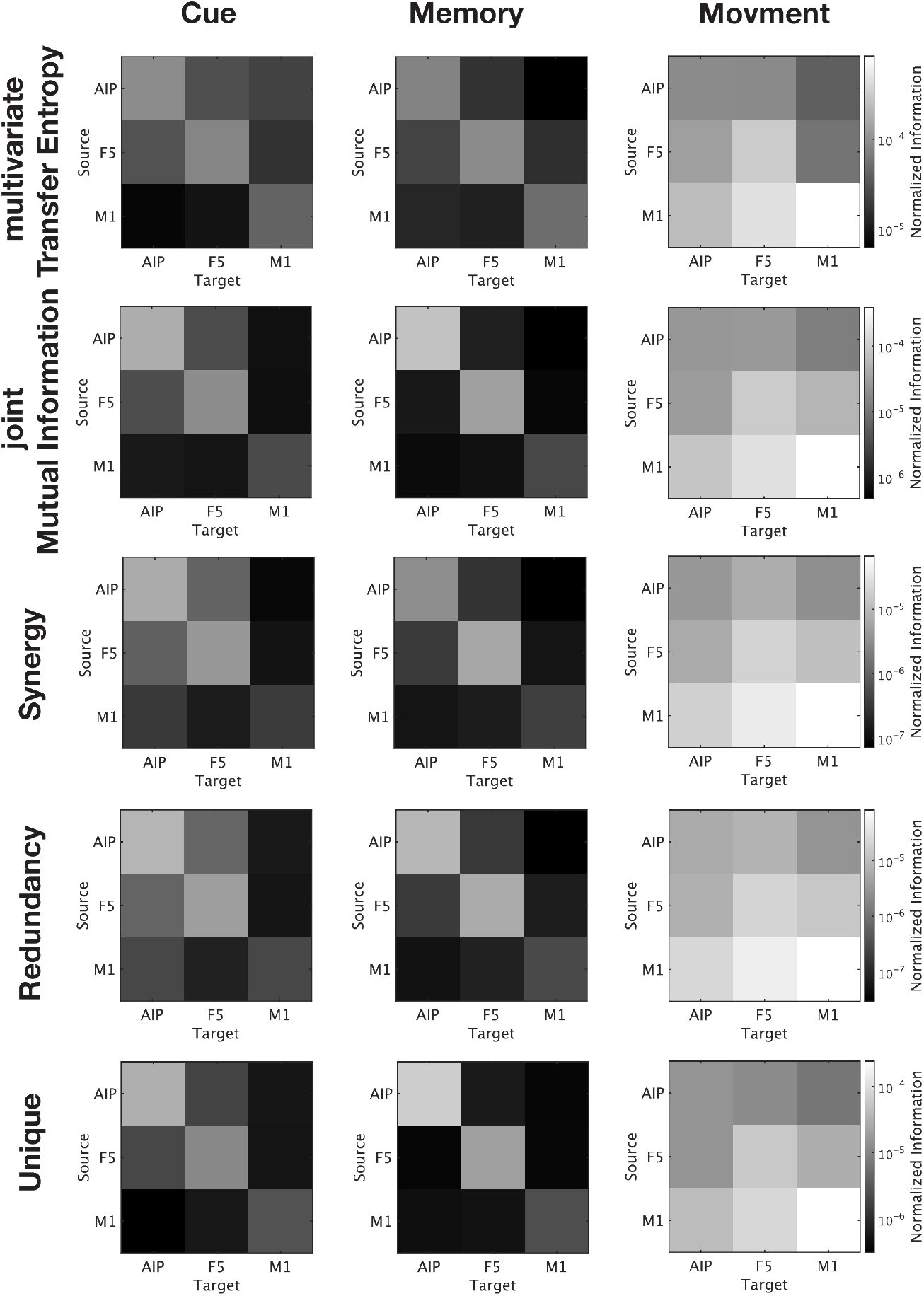
Changes in different types of Information processing within and between areas during different behavioral epoch. Heat maps representing the three normalized information dynamics (mTE, joint Mutual Information, synergy, redundancy, and unique) for three behavioral epochs (cue, memory, and movement), for each of the three areas (AIP, F5, M1)within and between areas in the feedforward and feedback direction. While the overall patterns of all information processing types are largely consistent, large differences between behavioral epochs are apparent. Moreover, during movement execution, information processing in the feedback direction is noticeably higher than in the feedforward direction between all areas.

Taken together, these results suggest that (1) processing between areas in the feedback direction is essential for this supposedly simple task and (2) that the extent to which information is processed between areas depends strongly on behavioral state, in contrast to information processing within areas.

#### 2.3.1 Network Dynamics

To assess how information processing changes over time and across conditions in fine detail, we next analyzed the information transfer network structure [41, 42]. By analyzing information transfer at the network-level with neuron resolution, we observed that the onset of the cue is associated with a transient reconfiguration of the network structure (see Fig. 6A-C). While the total amount of information transfer across the networks did not significantly change in response to cue onset, the normalized clustering coefficient, an indicator of local connection density, was significantly increased during the cue epoch for both grip-types A (precision grip) and B (power grip (cluster-based surrogate test, *p* < 10^−20^ for all windows and all grip types, see Methods) (See Sec. 4.7.3 [43, 28]. Similarly, for both grip types, the rich-club coefficient, an indicator of connection density of strongly connected neurons, was significantly higher during the cue epoch for both grip-types (cluster-based surrogate test, *p* = 0.002 for Grip A and *p* = 0.048 for Grip B). We can also see a significant increase in the rich club coefficient for both grip types during the movement epoch (*p* < 10^−20^ for all windows and all grip-types). Together, these results suggest that the transition from a passive cognitive state (fixation, memory) to an active cognitive state (perception of a visual cue or motor execution) is associated with marked changes to the information processing network structure. Furthermore, these changes are lost when considering average information dynamics of individual neurons (see Fig. 3 D-F for comparison). For example, an increase in the rich-club coefficient is seen in both the cue epoch and during the movement epoch, suggesting that the presence of rich-club structure is particularly important for active states. Interestingly, the emergence of a rich-club structure as well as an increased cluster coefficient occurs during the cue epoch without a corresponding change in the total amount of information transfer of the network, suggesting that it might be a true re-allocation of a fixed quantity of computational resources.

**Figure 6:**
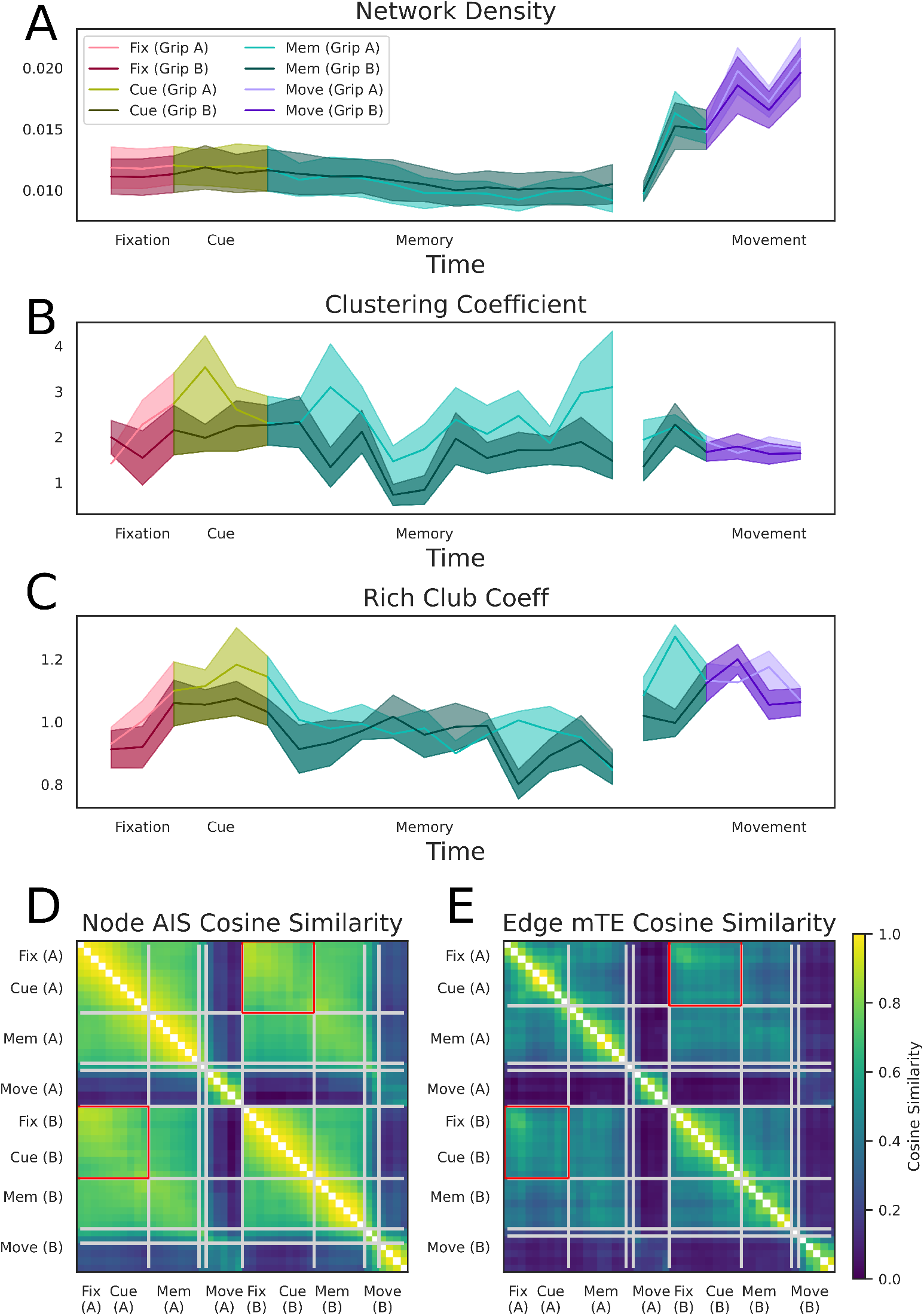
Differences in the effective network structure between behavioral states and conditions. **A-C** Window-level time-resolved time series for the three measures of network topology: network density (**A**), normalized global clustering coefficient (**B**), and the normalized rich club coefficient (**C**). The network density increased dramatically in the movement epoch but remained constant during all “cognitive” states. In contrast, the networks showed a transient reconfiguration during the cue epoch: increasing the hierarchical rich-club structure and becoming more clustered. **D-E** Pairwise cosine similarity matrices of AIS neuron-level and mTE connection-level values for over all time windows of both conditions. Grey lines indicate the boundaries of similar information network structure clusters as determined with a clustering algorithm. Note that the fixation and cue epoch clusters of both conditions belong to one cluster, indicated by the red box around the off-diagonal part of the cluster. Also note that the inferred clusters map onto boundaries between different behaviors.

**Figure 7:**
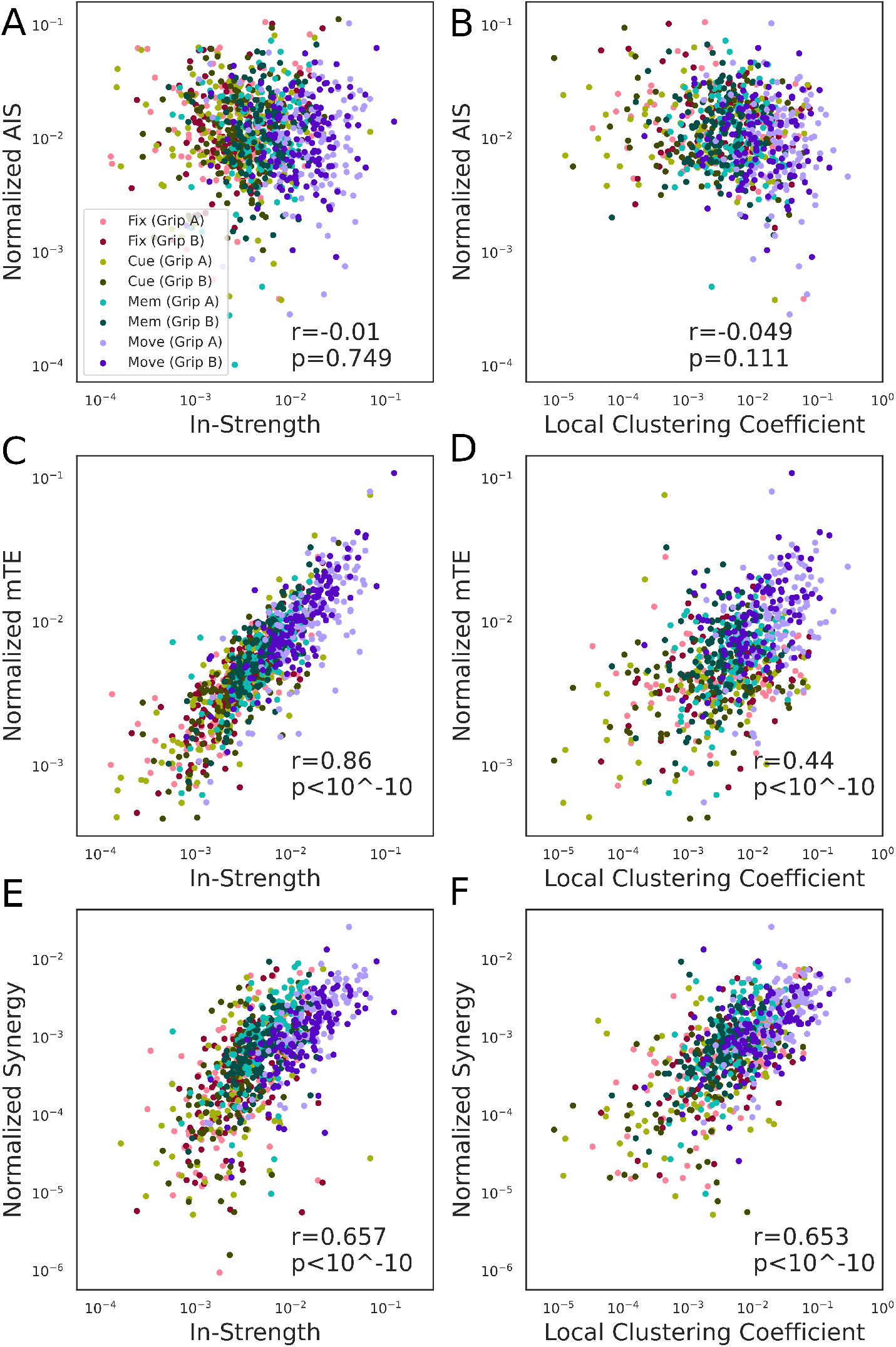
Information dynamics & local network structure. Plots showing how neuron-level information dynamics are related to neuron-level network measures (Spearmanm correlation coefficient). The color scheme is the same as in previous plots. **A-B** We found no significant relationship between active information storage and the in-strength of a neuron or the local clustering coefficient. **C** There was a highly significant correlation between the average transfer entropy of a neuron and its in-strength (*r* = 0.869, *p* < 10^−10^, which is unsurprising as both measures quantify the total information flow into a neuron, albeit in slightly different ways. **D** There was a significant correlation between average transfer entropy and local clustering coefficient (*r* = 0.44, *p* < 10^−10^), which shows that the local community structure a neuron is embedded in reflects the associated intensity of information flow. **E** The triadic synergy was positively correlated with the in-strength of target neuron (*r* = 0.657, *p* < 10^−10^), replicating the findings of [24]. **F** The triadic synergy was positively correlated with the local clustering coefficient (*r* = 0.653, *p* < 10^−10^), suggesting that extended local neighborhood structure plays a role in computation.

**Figure 8:**
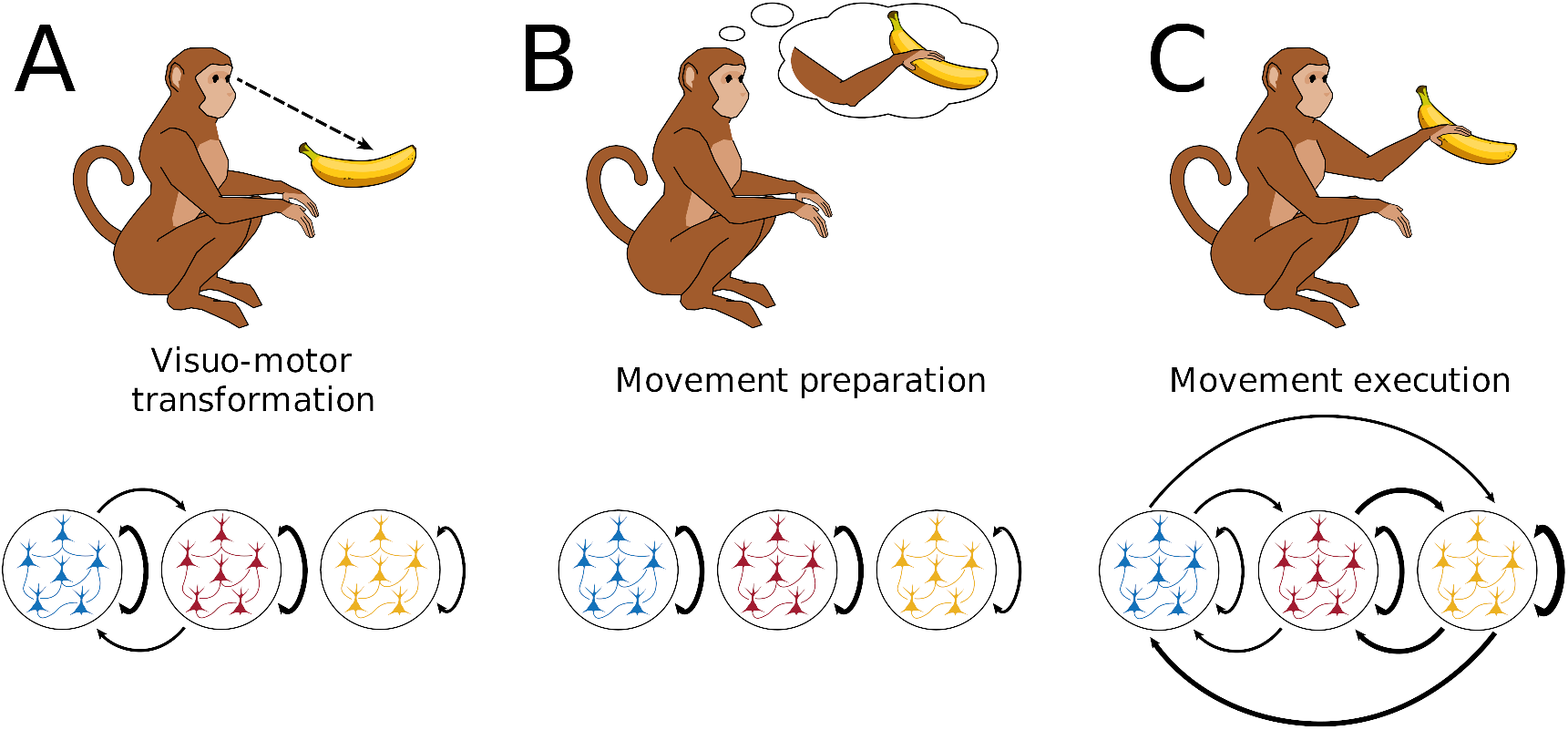
Variable feedforward and feedback dynamics during different behavioral states. Here we can see how different cognitive and behavioral epochs are associated with different patterns of information processing within and between areas. During the cue and early memory epoch, when monkeys are transforming the information from their environment into a movement plan, strong inter-area information processing is present between AIP and F5 in both the feedforward and feedback directions. In contrast, during the memory epoch where monkeys have to retain the instructed movement, information processing were primarily present within areas. Finally, during the execution of the movement information were processed globally within and between ares, outgoing from M1, with a dominance of processing in the feedback directions.

#### 2.3.2 AIS and mTE Similarity Clustering

In addition to various behavioral states monkeys also performed two different grasping conditions. Although all analyses above did not reveal any differences between the two grasp conditions (Figure 2, 3A-C), the possibility remains that differences in the information dynamics exist at the of fine-grained network-level with neuron resolution, which are lost at any larger scale. For this purpose, we examined condition dependent differences in the AIS values across neurons, and mTE values across network connections. For both measures, we estimated the cosine-similarity between the neurons and connections, respectively, of all time windows of all behavioral states and for both grip-types. The resulting cosine-similarities are depict as similarity matrices (see Fig. 6D-E). For both matrices, the values on the diagonal represent the similarity over time of first the precision grip (A) and then the power grip condition (B). The off-diagonal values thus represent the similarity between conditions over time. Over the time course of the task in both conditions, the AIS values across neurons and the mTE network structure were most dissimilar between the movement period and all other epochs. On closer examination, however, there are also differences between the fixation and cue epoch and the memory epoch. In direct comparison of both conditions, we can observe that the AIS values across neurons and the mTE-networks were most similar during the fixation epoch. Following fixation the similarity of the networks decreased, and condition-dependent difference became apparent during the cue, memory and movement epoch. Note that despite the similarity of network reconfiguration of both conditions over time, the fine-grained information dynamics at the network-level are highly dissimilar during the movement period. To verify behavioral state and condition differences, we used a multi-resolution consensus clustering (MRCC) algorithm [44] (see Sec. 4.8). MRCC is an unsupervised community detection algorithm that in this case identifies groups of AIS and mTE values over time and across conditions of increased similarity (see the squares along the diagonal in figure 5D,E). MRCC revealed that all but the combined fixation and cue epoch of both conditions (see the red squares in figure 5D,E) formed separate clusters. The across condition fixation and cue cluster is to be expected because the fixation epoch is the only epoch that is independent of the behavioral condition. Thus, this result confirms that the AIS values across neurons and the mTE network structure are (1) not statistically different during fixation and (2) statistically different during all other epochs. Note that the across condition cluster also comprises the cue epoch, which is probably due to the large sliding window size of 800ms. Interestingly, the independent condition specific clusters identified by the MRCC algorithm showed a similar temporal structure for both conditions resembling behavioral states such as memory and movement. Taken together, despite of no significant condition-dependent differences on average, both AIS and mTE showed significant differences at the neuron- and connection-level, respectively. Thus, these results suggest that the fine-grained structure of the information dynamic networks reconfigures for different conditions, forming distinct “information processing architectures”.

### 2.4 Relating Structure & Functional Information Dynamics

Previous studies performed on organotypic brain slices showed that synergistic information dynamics were highest between neurons with a high degree [24, 25, 26]. Yet, the meaning of synergistic processes in neural networks remains unclear. Therefore, it is important to scrutinize this relationship in the intact brain during behaviorally relevant processing. To assess this, we correlated each of the information dynamics for every neuron against local network measures (local clustering coefficient and in-strength, as a proxy for local rich club). We found no significant relationships between either in-strength or local clustering coefficient and node-level AIS. However, we found a strong, significant correlation between the in-strength of a neuron and the average mTE flowing into it (r=0.86, p< 10^−10^). While this is unsurprising, it is not entirely trivial: a low in-degree neuron and a high in-degree node might have the same in-strength, but different average incoming transfer entropies. Similarly, we found a strong, significant correlation between the in-strength of a neuron and its local clustering coefficient (r=0.44, p< 10^−10^), which indicates that the information processing occurring in a single neuron is informed by the local connectivity pattern of that neuron’s neighbors. Finally, we found strong, significant correlations between synergy and both in-strengths (r=0.657, p< 10^−10^) and local clustering coefficient (r=0.653, p< 10^−10^). As with the transfer entropy results, these results show that a neuron’s location in the effective connectivity network relative to other neurons, informs the type of information processing it predominately performs.

## 3 Discussion

In this work, we have used information theory to study behavior-related changes in patterns of information processing in neural networks spanning multiple brain areas of the macaque fronto-parietal grasping network. We investigated (1) how information processing changes during different cognitive and behavioral states over the course of a behavioral task (2) to what degree information is processed restricted to within specific areas, or distributed between areas, (3) to what degree this inter-area processing structure is fixed or changes dynamically for different behavioral states, and (4) whether behavioral condition-dependent differences are present at the fine-grained information processing network structure. For this purpose, we estimated different types or components of “information processing” (referred to as “information dynamics” [45]): the active information storage (how the past activity of a single neuron informs on its future), transfer entropy (how the past of a set of source neurons informs on a single target neuron’s future), and the information modification/synergy (information from the joint-state of multiple sources about a target’s future that is irreducible to any simpler combination of sources).

We found that information processing of different types changes during different cognitive and behavioral states. Furthermore, the degree of information processing taking place within a single area, as opposed to between areas, strongly depended on the behavioral state. During visuo-motor transformation area AIP and F5 formed a processing unit (Figure 8A). During memory, this unit unexpectedly fell apart, despite the amount of information processed *within* each area remaining constant (Figure 8B). Finally, the highest level of information processing occurred during the movement execution. In contrast to the two previous behavioral states, movement execution appeared to be a global process with the greatest amount of computation occurring between all areas, especially M1 (Figure 8C). Movement execution-related processing between areas was more pronounced in the feedback direction suggesting that online feedback to earlier areas is an essential part of movement execution. While the the average information processing dynamics were the same for different conditions across the different cognitive and behavioral states, significant condition-dependent differences became apparent at the fine-scale network level with neuron resolution. Our results suggest that connected areas can dynamically form functional units that together enable the required cognitive or behavioral state, while fine-grained reconfigurations of the network structure reflect different behavioral conditions. These ensembles of areas are flexible: multi-area units can subsequently fall apart to reform into other multi-area processing unit according to the cognitive and behavioral demands.

This work unifies previous findings about information dynamics, behavior-related neural population dynamics, and the structure of networks of neurons Prior work on information dynamics has demonstrated the existence of synergistic processing in dissociated cultures [20, 25]: synergistic dynamics can be sensitive to local network structure [36], and vary over time [21]. Similar analyses have been done for AIS [15] and TE [16, 17], although until now it has been a mystery how these different modes of processing relate to cognition and behavior.

Previous work on fronto-parietal neural rate dynamics has demonstrated that neural populations respond to movement preparation as well as movement execution, but with independent dynamics for both processes [37, 46, 47]. Therefore, preparation-related information must be “transformed” into movement execution-related information. However, analyses of population dynamics do not allow a direct estimate of computation. Historically, the amount of information processed was either estimated based on correlation analyses [48] or indirectly inferred via an artificial neural network model [49]. Prior work on the inference of network connectivity at the neuron-level has demonstrated the presence of strong inter- and intra-are connectivity and an area spanning rich-club of neurons [28], but its behavioral relevance was elusive.

We found significant cognitive and behavioral state-related changes to information processing architecture. mTE and synergy both significantly increased in the movement epoch, while synergy additionally increased during the cue epoch (Figure 3). The observed increase in information processing in both epochs is in agreement with the cognitive and behavioral requirements: during the cue epoch, monkeys must actively observe and integrate instructions, while during the movement epoch, monkeys must actively move arm and hand to grasp the target with the required grip type. In contrast to these active periods, monkeys must remain still during fixation and memory epochs, which suggests a lower degree of information processing during these steady periods. This assumed relationship between the estimated amount of information processing and the behavioral requirements is further supported by the finding that the mTE effective network dynamics vary between active and steady states (Figure 6A-C). In particular, the presence of an effective rich-club network structure distinguished active and steady states (no significant rich-club during steady states, significant rich-club during active states).

Not only did information network dynamics resemble the cognitive and behavioral states, but the increase in information processing and network dynamics preceded the movement onset. The so-called internal movement onset in the brain, however, has been shown to precede the real movement onset [37, 38] simply because nerve signals take time to arrive at the appropriate arm muscles and arm muscle potentials need time to build up to lift of the arm and hand. Therefore, the observed mismatch of information processing with the movement epoch also further supports the assumed relationship between information dynamics and behavior. Although the lack of significant differences in average information processing between conditions seems disappointing at first glance, it further validates our findings. Both grasp conditions have equally salient cues, can be assumed to be equally difficult to prepare for, and presumably requires a comparable amount of effort to perform. Given these facts, it is not surprising that the total amount of information required for both was similar. Similar level of information processing between the two conditions serves as cross-validation of our results. Thus, our findings suggest that the estimated information dynamics at the network-level capture the true behaviorally relevant processing or at least parts of it.

Given the evidence that information theory allows us to capture the underlying processing in the fronto-parietal network of neurons, three questions arise: (1) Why is information processed between areas and not exclusively within areas? (2) Why do the areas that form a processing unit change depending on the behavioral state? (3) How are different multi-area processing units formed?

Regarding the first question, there exist compelling evidence that large populations of neurons in multiple areas are modulated in a similar way during the same cognitive or behavioral task [1, 2, 3, 4]. This high degree of correlated activity or information is made possible by the strongly reciprocal connection of mammalian cortical areas [50, 51]. Additionally, there is strong evidence coming from neuroimaging that the pattern of correlated activity is constantly changing despite static connectivity [52, 53]. These results suggest that the cortex should be viewed more as a single interconnected network of neurons than as a structure divided into areas. Consequently, information is likely to be globally communicated and presumably also processed. Our results of strong processing in both the feedforward and feedback directions provide evidence for this proposal. A more global integration may also allow for a greater flexibility and the integration of more factors simultaneously than a classical hierarchical processing structure.

Regarding the second question, it may be beneficial to flexibly recombine processing from multiple areas according to behavioral demand. For the given task, first visual information must be transformed into a movement plan. It has been demonstrated that areas AIP and F5 are involved in visuo-motor transformation, but M1 is not [5, 6, 8]. Therefore, it makes sense that AIP and F5 form a processing unit during visuo-motor transformation not involving M1. Neurons of both areas also show elevated and prolonged activity during memory periods [5, 6]. Surprisingly, we found that the processing units of AIP and F5 fell apart during memory, although memory may require less “processing” and something more akin to “storage” (not to be confused with the measure of AIS reported here). A large number of studies have shown that mainly M1, but also F5, are involved in movement execution [37, 54, 55, 8]. In recent literature, a mechanism has been proposed for how a movement plan is translated into movement execution in the premotor and motor cortex [56]. Our finding of high levels of information processing within and between M1 and F5 during movement execution is consistent with this hypothesis. In contrast, little is known about the involvement of area AIP in movement execution. Only a single study showed that the movement trajectories from the whole arm and hand can be decoded from AIP during movement execution [7]. Thus, our results suggest that the number of cortical areas involved in the required processing for movement execution is larger than previously thought. We speculate that the strong feedback processing might be necessary to align precise details of the movement with different sensory areas, the evaluation of the movement, or learning.

Regarding the third question, a rich literature has explored how dynamic communication is established between cortical areas [57]. However, most results about dynamic communication capture co-fluctuations of different brain signals and reveal little about multi-area processing units. Nevertheless, oscillatory synchrony has been proposed as a coordinative mechanism for inter area communication [28], which might also be involved in the dynamic formation of multi-area processing units.

Another intriguing finding is that condition-dependent differences of the fine-scale information processing network structure were present during visuo-motor transformation, memory and movement execution, despite of no significant differences of the average amount of information processing during all three cognitive and behavioral states (Figure 3D-F; 4; 6D,E). As mentioned before, both grasp conditions have equally salient cues, can be assumed to be equally difficult to prepare and are require a comparable level of effort to perform. Therefore, a similar average level of information processing may be expected. On the other hand, it is also evident that information processing must be different for the two grasping conditions. Consequently, the observed differences in the processing network structure at the neuron-level most likely reflect the distinct, fine-scale neuronal computations underlying different conditions. Thus, the methods described here provided a powerful tool to study neural computations in the network during different cognitive and behavioral processes. In this first study examining the behavioral state and condition dependent differences in the information processing network structure, it becomes apparent that behavioral state differences are larger than condition dependent differences. Moreover, the network state during fixation as well as during cue and memory of both condition is more similar to each other than the network state during movement execution of both conditions. We suggest that the active execution of a movement leads to a much more drastic change in the information processing structure than different cognitive states. In contrast, the finer but distinct differences between conditions remained relatively stable. Taken together, these results suggest a hierarchy in information processing in which the coarse network structure determines the behavioral state and finer changes in the network structure reflect different conditions.

This study has some limitations that should be discussed. One of the most significant limitations is that, while multivariate transfer entropy gives a much more “complete” picture of the information transfer dynamics (and appears to better estimate the generative network [33]), it suffers from a terrible curse of dimensionality and will not scale gracefully as the number of neurons being recorded from gets large and/or the number of bins of history increases. Consequently, for researchers interested in the role of redundancies and synergies in time series dynamics, heuristics such as the O-information may be useful [58, 59, 60]. The partial information decomposition (PID) framework suffers from a similar curse of dimensionality as the the mTE algorithm: the number of unique partial information atoms grows super-exponentially with the number of sources. Since neurons can participate in hundreds or thousands of synaptic relationships, being restricted to sets of 1-5 interacting neurons is a severe limitation. Consequently, there is an urgent need for analytic techniques to identify synergistic relationships that do not require brute-forcing the entire partial information lattice. There is also a long-standing question about how best to address the issue of variable firing rates in neural recordings. One could argue that firing rate represents a confound that must be “conditioned out”: for example, a higher-firing rate, but random, pair of neurons will likely have greater apparent transfer entropy relative to a lower firing-rate pair, simply because there is more entropy in the system overall, even if no relationship actually exists [17]. Alternatively one could argue that the firing rate *is* the channel: changing the level of neuronal activity appears to be at least one way that neurons communicate information (known as “rate coding”, evidence for which dates back to the early days of neuroscience [61]). Alternative coding schema (such as temporal coding [62]) which rely on relative timings between neurons still can make use of changes in firing rate since higher entropy processes can support a wider range of firing rate patterns, including relatively low-probability patterns. The question of how best to, or whether to control for firing rate at all becomes ambiguous, especially since the exact nature of the neural code remains mysterious and likely combines many different dimensions including rates, spiking timing dependent codes, and more [63, 64]. This study demonstrates two fairly liberal normalizations: for the individual information dynamics results (AIS, mTE, synergy), we normalized the raw information values by the entropy of the receiver neuron to approximate the proportion of uncertainty explained by each mode. For sparse processes, this serves as a simple firing-rate correction, since low-firing rate neurons will have low entropy and vice versa for relatively high-firing rate neurons. Crucially, this simple correction is easily interpretable, it only requires information-theoretic measures (keeping the analysis conceptually simple), and computationally practical (not requiring huge numbers of null models as in the AIS and mTE inference). The second normalization is done in the redundancy/synergy bias calculation, where each partial information atom is normalized by the total mutual information *I*(*S*_1_, *S*_2_; *T*). This provides a different interpretation of the information: rather than uncertainty resolved, it is the proportion of the total information flow that is shared or synergistic, and not specific to any single source or target neuron. These different analytic choices throw different statistical relationships into the spotlight and highlight how it can be limiting to consider only one “true” correction and instead keep the pipeline flexible.

In total, information dynamics represents an appealing framework with which to explore the brain as an integrated whole, and can be easily adapted to assess computational processes at multiple scales. Given the extensive public data available to neuroscientists, we are optimistic that information dynamics can provide new insights into the relationship between brain, behavior, computation, and dynamics. In particular, information dynamics marries two well-established frameworks in modern neuroscience: computational approaches, and dynamical systems approaches [65, 66, 38, 67, 49, 56]. Sometimes called “computational mechanics” [68], information dynamics analysis can provide a bridge between approaches to provide insights that may not be obvious from either one.

## Author Contributions

TFV, OS, and BD collectively conceptualized the project. HS and BD prepared experiments. BD and SS collected and preprocessed the data. TFV, OS, and BD performed analysis of the data. TFV and BD wrote the initial draft of the manuscript. TFV, OS, HS and BD provided feedback and editing of the manuscript. All authors approved final submission.

## Acknowledgements

TFV is supported by NSF-NRT grant 1735095, Interdisciplinary Training in Complex Networks and Systems. TFV and OS are supported by NIH R01 MH121978. BD and HS are supported by the German Ministry of Education and Research (FKZ 01GQ1903). We would like to thank Natalie Bobb, Ricarda Lbik, and Matthias Dörge for their assistance with animal handling and setup maintenance. Moreover, we would like to thank Dr. Joe Lizier for advice throughout the process, particularly in regards to the use of the IDTxl package, as well as Dr. Patricia Wollstadt for her help as the maintainer of the IDTxl repository, and all of the other developers who contributed. Furthermore, we would like to thank Dr. Samantha Sherrill for her insight and experience using PID in the context of single neuron recordings, as well as Drs. John Beggs and Ehren Newman and their respective labs for insightful feedback.

## 4 Materials & Methods

### 4.1 Basic procedures

Neural activity was recorded simultaneously from many channels in two female and one male rhesus macaque monkeys (Animals S, Z, and M; body weight 9, 7, and 10 kg, respectively). Detailed experimental procedures have been described previously [69, 28]. All procedures and animal care were in accordance with German and European law and were in agreement with the Guidelines for the Care and Use of Mammals in Neuroscience and Behavioral Research (National Research Council, 2003)

### 4.2 Behavioral Task

Figure 2 illustrates the time course of the behavioral task described in detail previously [69, 28]. Monkeys were trained to perform a mixed free-choice and instructed delayed grasping task, with the exception of monkey M, which was trained to perform only a instructed delayed grasping task. They were seated in front of a handle that could be grasped in two different ways and visual cues were displayed on a masked monitor that was superimposed on the handle using a beam splitter mirror. Trials started after the monkey placed both hands (for monkey S) or one hand (for monkey Z and M) on the resting positions. For monkey Z and M, the non-task relevant arm was comfortably restrained in an acrylic glass tube during the experiment. Next, a small red disk was displayed centrally on the monitor, which had to be fixated for a variable period of 600 to 1000ms (fixation epoch). After the fixation epoch, cues in the form of larger disks were shown next to the fixation disk for 300ms and the handle was illuminated. In the instructed context, one of two discs was displayed instructing the monkey about the required grip type (power or precision; cue epoch). In the free-choice context both disks were displayed indicating the monkey to freely choose between the two grip types. After the cue was turned off, the monkey had to remain steady for a variable time of 1100 to 1500ms (memory epoch). The turning off of the central fixation disk instructed the monkey to reach and grasp the target with the required grip type (movement epoch) to receive a liquid reward. Note that to encourage the monkey to perform both grip types during the free-choice context, the reward was iteratively reduced every time the monkey repeatedly chose the same grip type (fraction of power grip selections: 38.5±4.2% SD and 53.3±5.2% SD for S and Z, respectively; Note that no choice ratio is available for monkey M, since he performed only instructed trails). All trials were randomly interleaved and executed in darkness. Note that the behavioral task also contained delayed instructed trials for monkey S and Z, which were not analyzed in this study. To increase the reliability of the information dynamics estimation, instructed and free choice trials for the same grip type were pooled for all further analyses.

### 4.3 Electrode Implantation and Data Recordings

Surgical procedures have been described in detail previously [69, 28]. Briefly, two floating microelectrode arrays (FMAs; Microprobes for Life Sciences; 32 electrodes; spacing between electrodes: 400*μ*m; length: 1.5 to 7.1 mm monotonically increasing to target grey matter along the sulcus) were implanted per area in the ventral premotor cortex (area F5), in the anterior intraparietal area (AIP), and in monkey M also in the primary motor cortex (M1) for a total of 128 electrodes for monkey S and Z, and 192 electrodes for monkey M. Monkeys S and M were implanted in the left and monkey Z and the right hemispheres (Figure 1B).

Extracellular signals from the implanted arrays were amplified and digitally stored using a 128-channel recording system for monkey S and Z (Cerebus, Blackrock Microsystems, sampling rate 30 kS/s, 0.6–7500Hz band-pass hardware filter) and a 256 channel Tucker-Davis system for monkey M (TDT RZ2; sampling rate 24.414 kS/s, 0.6-10,000 Hz band-pass hardware filter) while the monkeys performed the delayed grasping task. All data were saved to disk and analyzed using custom MATLAB code (The Mathworks Inc., Natick, MA).

### 4.4 Data Preprocessing

For spike detection, the broadband signals were first low-pass filtered with a median filter (window length 3ms) and the result subtracted from the raw signal, corresponding to a nonlinear high-pass filter. The signal was then low-pass filtered (4th order non-causal Butterworth filter, fc: 5000 Hz). To eliminate common noise sources, principal component (PC) artifact cancellation was applied for all electrodes of each array, as described previously [70, 28]. To ensure that no true spikes were eliminated from individual channels, PCs with any normalized coefficient greater than 0.36 (conservatively chosen) were retained. Spike waveforms were detected and semi-automatically sorted using a modified version of the offline spike sorter WaveClus [71, 72, 28].

Units were classified as single- or non-single unit based on five criteria:

1. The presence of short (1–2ms) intervals in the inter-spike interval histogram for single units;
2. The similarity of all waveforms of each unit at each time point;
3. The separation of waveform clusters in the projection of the first 13 features (a combination for optimal discriminability of PCs, single values of the wavelet decomposition, and samples of spike waveforms) detected by WaveClus;
4. The presence of well-known waveform shapes characteristics for single units;
5. The shape of the inter-spike interval distribution.

After the semiautomatic sorting process, redetection of the different average waveforms (templates) was done to detect overlaid waveforms [73, 28]. To achieve this, filtered signals were convolved with the templates starting with the biggest waveform. Independently for each template, redetection and resorting was run automatically using a 32-dimensional linear discriminate analysis for classification of waveforms. After spike identification, the target template was temporally matched and subtracted from the filtered signal of the corresponding channel to reduce artifacts during the detection of the next template. This procedure allowed us to detect spikes with a temporal overlap up to 0.2ms. Unit isolation was evaluated again, based on the five criteria mentioned above, to determine the final classification of all units into single or non-single units. Stationarity of firing rate was checked for all units and in case it was not stable over the entire recording session (more than 30% change in firing rate between the first 10 min and the last 10 min of recording) the unit was excluded from further analyses (< 3% of all single units). Note that only well isolated single neurons based on the five criteria above were used for all further analysis.

### 4.5 Sliding Window Analysis

To assess how effective connectivity patterns change with the behavior of the macaque, we made use of a sliding-windows analysis [74]. We used 800 ms windows, incremented with 100 ms tied to either the onset of the cue condition (starting with −800ms from the onset of the cue and continuing for 1700 ms), or the onset of movement (starting with −800 ms from the onset of movement and continuing for 600 ms.) The result was ≈900 individual slices across the 10 different recordings.

For each of these slices, we inferred a single multivariate transfer entropy network using the IDTxl package [34], as well as the local active information storage, and triadic synergies following the pipeline outlined in [24, 25].

We chose the sliding window approach to temporal localization instead of the more fine-grained analysis of local information dynamics [75] to ensure that we could see how the overall network topology changed in time: a local information approach would have improved our temporal resolution but at the cost of providing a static network, since fixation, cue, and movement conditions would need to be aggregated to construct a single joint probability distribution (which the system presumably visits different parts of in different conditions, which the local analysis would reveal). This would have precluded a dynamic network approach, which is a core aspect of this study.

### 4.6 Active Information Storage

Active information storage (*AIS*) is arguably the simplest of the three information dynamics explored here, and quantifies how much information about the next state of the neuron is encoded in the past *k* states [75]. *AIS* can be thought of as a measure of the “memory” capacity of the neuron: if every subsequent state is chosen at random, then *AIS* = 0, as knowing the past does nothing to reduce our uncertainty about the mediate future, however if the past states of the neuron affects the probability of a particular next state, then the system can be thought of as “remembering” that information and “using” it when deciding what the next state will be (note the scare quotes). A simple example might be the refractory period that follows an action potential: following an action potential, most neurons cannot immediately fire again, as the correct charge needs to be built up. Consequently, an observer attempting to model the dynamics of a single neuron would know that the probability of a spike goes down if the neuron is known to have fired in some previous time-window, and this would be reflected in the *AIS* measure. For a more involved discussion of *AIS* in understanding neural dynamics see [15].

The *AIS* of a single variable *X* at time *t* is given as the mutual information between the immediate state of *X* at time *t* and the embedded joint-state of the past k time-steps (the embedding here is similar to a Takens embedding [76]):

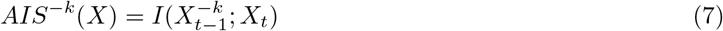

Intuitively, this can be understood as measuring how much knowledge of the joint past reduces our uncertainty about the immediate future. The free parameter *k* indicates how many steps into the past we want to consider when calculating the *AIS* and should be chosen with some care. If *k* is too small, then we run the risk of missing out on relevant information from more than *k* steps in the past and we will under-estimate the true *AIS*. On the other hand, if *k* is too large, then we run the risk of under-sampling the joint-probability space and over-estimating the true *AIS*.

In this work, we set *max k=3*, which given 5 ms bins works out to a considered history of 15 ms (based on prior work reported in [77, 78]), and then used the IDTxl package [34] to infer an optimal non-uniform embedding. Following [25, 26], the AIS of a neuron was normalized by dividing it by the total Shannon entropy of the neuron. This provides a simple correction for variable firing rates, removing the confound that higher-entropy neurons have a greater capacity for any information dynamic.

### 4.7 Multivariate Transfer Entropy Networks

While the *AIS* is restricted to understanding the information dynamics of a single neuron (how it is “remembered” through time), the transfer entropy (*TE*) describes how information “flows” from one source neuron (or a set of source neurons) to a single target neuron [75, 27, 79] For a source *X* and a target *Y, TE*(*X* → *Y*) is given by the mutual information between *X*’s past and *Y*’s immediate next state, conditioned on *Y*’s own past.

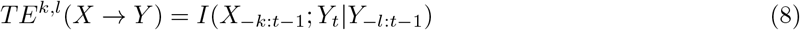

By conditioning on *Y*’s past, we are able to extract the information that *X*’s past provides about *Y*’s future *above and beyond the AIS in Y*. Transfer entropy has become an established tool in modeling neural data [80, 16, 29, 24, 59]. A common technique is to use transfer entropy to construct an effective connectivity graph meant to model the directed synaptic structure of the underlying biological network, to reveal properties like rich clubs [81, 25], multiplex structures [82], and communities [83].

The “classical” transfer entropy has a significant limitation, however that it is strictly bivariate, and consequently blind to higher-order information dynamics. Consider the case of two source neurons *A, B* synapsing onto a single single target neuron *C*, which implements *C_t_* = *XOR*(*A*_*t*−1_, *B*_*t*−1_). Due to the synergistic nature of the logical XOR, bivariate transfer entropy will not find any relationship between *A, B* and *C*, since information about *C_t_* is only disclosed by the joint pasts of *A* and *B* considered together. To address this, we can use the conditional transfer entropy, which quantifies the information flow from one neuron to another *in the context of other neurons in the system*. Formally:

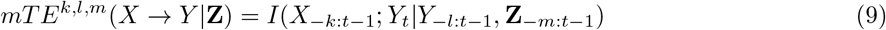

Where **Z** can be a multidimensional variable. Note again that each variable can have a different lag, to account for differences in the intrinsic dynamics in each element. In our simple logical XOR example, *TE*(*A* → *C*) = 0 bit, but *TE*(*A* → *C*|*B*) = 1 bit.

For a would-be network neuroscientist, this presents an optimization problem: for a pair of neurons *X* and *Y*, how do we find the smallest set of neurons *Z* such that the context *Z* provides discloses *all* the relevant information flow from *X* to *Y*. The answer is the multivariate transfer entropy (*mTE* inference, detailed in [32], which uses greedy optimization and multi-level statistical hypothesis testing to define a “parent” set for every neuron in the network, from which the optimal conditional transfer entropy between every parent in the parent set and the target neuron can be derived. Work using simulated data has shown that the bivariate and multivariate transfer entropies return networks with strikingly different topologies and that the multivariate transfer entropy networks are reliably closer to the “ground truth” [33, 84]. It has been established previously that non-trivial synergistic information dynamics exist in biological neural networks [25, 85, 26], suggesting that bTE, which is known to be blind to synergistic relationships, is missing important aspects of the system’s dynamics.

As with the *AIS*, the choice of embedding parameters is important, and to maintain consistency, we again set the maximal *k* value to 3, and used the IDTxl package [34] to find optimal non-uniform embeddings for each neuron. Due to the large number of neurons recorded, we inferred the parent set of each neuron independently and constructed a putative *mTE* network post-hoc without correcting for familywise error rates at the whole-network level (every edge was corrected for multiple comparisons). This is an acknowledged limitation of our inference, however the computational requirements for a full, error-corrected inference are beyond our capabilities at present. It may be possible that a subset of edges inferred are false-positives, although we do not anticipate that enough false positives would have passed the otherwise stringent threshold to substantively compromise the overall analysis.

#### 4.7.1 Density, Rich Club Coefficient & Clustering Coefficient

For a directed network *G* = (*V,E*), where *V* is the vertex set and *E* is the edge set, the density of the network is given as:

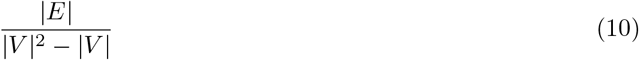

Which records the ratio of the number of edges that exist in the network to the total number of edges that could exist after self-loops are removed.

For a directed, weighted graph, the local clustering coefficient for a node *v* is given by:

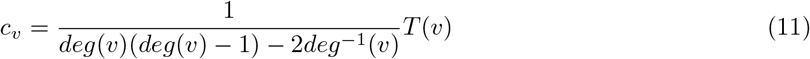

where *deg*(*v*) is the sum of the in- and out-degrees of *v*, *deg*^−1^ is the reciprocal of the degree of *v*, and *T*(*v*) counts the number of directed triangles that v participates in. We used the Networkx clustering toolbox [86] for this calculation. The average clustering coefficient for the whole network is given by:

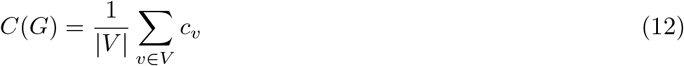

Which is simply the unweighted average of each local clustering coefficient.

The rich club coefficient quantifies the extent to which nodes of a network with a high number of connections are more strongly connected with each other then expected by chance [87, 88]. We calculated a modified version of the original weighted rich club coefficient as described in detail in the following [25]. The weighted rich club coefficient was calculated separately for all effective mTV networks (23 time windows x 2 grasping conditions). First, all neurons per networks were sorted by the combined in and out degree to obtain a degree rank *r_deg_* per neuron. Second, the weighted rich club coefficient Φ_*w*_(*r_deg_*) was consecutively calculated given by:

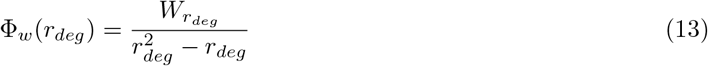

with *W_r_deg__* denoting the sum of weighted connections of the subset of neurons with a degree rank > *r_deg_*, and with 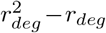 denoting the maximum possible number of connections of the same subset. Since neurons with a high degree have an increased probability by chance to be connected to other neurons with a high degree, the weighted rich club coefficient needs to be normalized by surrogate data based on networks with the same degree and ideally strength distribution. The estimation of the surrogate networks is described in detail below. In a third step, each rich club value was normalized by the corresponding average of 1000 surrogate weighted rich club coefficients. Finally, the normalized weighted rich coefficients were resampled to 1 to 100 using linear interpolation to average across different recording sessions with different numbers of neurons.

#### 4.7.2 Surrogate Networks

To estimate the normalized cluster coefficient, the normalized rich club coefficient and for statistical purposes we generated 1000 partitions of surrogate networks for each inferred multivariate transfer entropy network (21 time windows × 2 conditions × 10 recording sessions). Surrogate networks were generated by first shuffling network connections and second reassigning connection weights. Networks of neurons estimated from simultaneous extracellular recordings are constrained by multiple anatomical and technical factors. To ensure that the above network analyses and statistics are not biased by these factors, network properties affected by these factors should be preserved in the surrogate networks. Several studies have shown, that the number and strength of neuronal connectivity decreases with spatial distance [89, 90, 28]. Additionally, the spatial electrode configuration on each array as well as the configuration of the recording arrays to each other results in a spatial inhomogeneous subsampling of neurons [91, 28]. To account for both, we held the number of connections on the same electrode, the same array, the same cortical area, and the different inter-area connections the same and matched the average connection strength per distance category as close as possible. Cortical neurons also have an increased probability to be reciprocally connected to each other [92]. For this reason, we also held the ratio of reciprocal to unidirectional connections per distance category the same. Furthermore, for the normalization of the rich club coefficient it is necessary that the number of connections per neuron is preserved because strongly connected neurons have an increased probability of being connected to each other by chance [87, 88]. However, the same is true for the connection strength per neuron, which has been ignored in the literature so far. We therefore not only held the number of connections per neuron the same, but also matched the connection strength per neuron as close as possible. To our knowledge the described above surrogate network method is more conservative than any other method used in the field, which emphasizes the validity of the results based on this surrogate method.

#### 4.7.3 Cluster-Based Significance Testing

For the statistical testing of increased rich club and clustering coefficient over the time course of the task, we performed cluster-based surrogate tests. Statistical testing was performed across recording sessions and separate for the power and precision condition (Fig. 2 A) [43, 28]. The testing procedure was the same in both cases and the same set of surrogate networks was used as described above. The computation of the cluster-based surrogate tests was performed as following:

1. Test values (rich club coefficient or rich club coefficient) were calculated separately for all surrogate networks of all time windows and recording sessions.
2. Normalize all surrogate test values by dividing the average surrogate test values of the corresponding time window and recording session (normalized rich club coefficient or normalized rich club coefficient).
3. Average all surrogate test values per partition and time window (for the normalized rich club coefficient the interval between the 83rd and 95th percentiles) to obtain one value per network.
4. Calculate the surrogate t-values across recording sessions for all partitions and time windows.
5. Select all surrogate t-values larger than a thresholding criterion and cluster and sum them on the basis of temporal adjacency.
6. Take the largest summed surrogate t-value per partition to construct a distribution of largest summed surrogate t-values.
7. Repeat step 1-5 for the recorded test values.
8. For every summed t-value calculate the proportion of surrogate t-values that are larger than the recorded summed t-values, which corresponds to the p-value.
9. Compare each p-value with a critical alpha-level (0.05 in all cases).

Note that this single comparison replaces the multiple comparisons of the test-values over time.

### 4.8 mTE and AIS Distribution Similarity Analysis

Similarity matrices derived from the time-resolved TE and AIS analysis underwent clustering and community detection to extract, in a purely data-driven way, time periods that had a distinct TE similarity structure. To create similarity matrices for the distribution of AIS values across the nodes, we created a matrix where cell *ij* gives the cosine similarity between the vector of AIS values across all nodes in the *i^th^* and *j^th^* windows. Cosine similarity is given as:

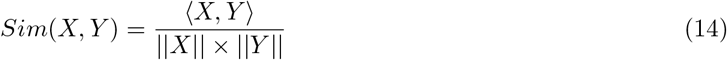

Where 〈*X, Y*〉 is the dot product of *X* and *Y*, and ∥*X*∥ is the norm of X. The implementation we used was from the Scikit-Learn Pairwise Comparisons package.

The mTE pipeline was the same, although we took the cosine similarity of the flattened adjacency matrices. First, similarity matrices were aggregated across the 10 trial repetitions by taking the mean. This resulted in a single 46×46 element matrix for both mTE and AIS. These matrices were then clustered using a version of modularity maximization (employing the Potts null model) and multi-resolution consensus clustering [44]. The resolution parameter, used by the Louvain algorithm, was stepped through a range of 1,000 values covering modular partitions yielding between 2 and 46 modules (the minimal and maximal number of modules possible). The resulting 1,000 partitions were aggregated into a single co-classification matrix which was scaled between followed by subtraction of an analytic null that captures the expected co-classification (mean propensity for each node pair to be grouped in the same community) under random permutation of the module assignments [44]. The resulting scaled co-classification matrices were then subjected to consensus clustering with *τ* = 0. The level of *τ* corresponds to the (constant) null. The resulting consensus communities correspond to time steps that are clustered together based on the similarity of their mTE/AIS similarities. Variations in the number of samples (between 100 and 10,000) and range of resolution parameter (2-23; 2-10) had no effect on the cluster boundaries.

### 4.9 Synergy & Partial Information Decomposition

The final information dynamic is *information modification*, which describes information that is somehow non-trivially altered by the interaction of two or more incoming “information streams” [75]. Information modification has historically resisted formalization, although one appealing proposal suggests that synergistic information could serve as a useful foundation with which to understanding information modification [35]. Recall that synergistic information is inherently multidimensional information that is not trivially reducible to any of its component parts: it represents information that is, in some sense, “greater than the sum of its parts.” Since its initial proposal, synergy has been used to great effect as a measure of non-trivial computational processes in neural cultures [25, 85, 26].

Calculating the amount of synergy present in dynamic information is a difficult task. The key development by Williams and Beer [39, 40] was that the joint mutual information between a set of variables and a single target could be decomposed into “partial information atoms”, which describes the information specifically encoded in a particular combination of source elements and none simpler (for an alternative derivation of the basics of partial information decomposition that leads to the same results, see [93]). For the case of two source variables *X*_1_ and *X*_2_ and a target *Y*, this leads to decomposition into redundant, unique, and synergistic partial information atoms:

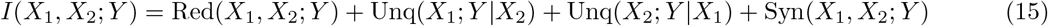

Where Red() indicates the information about Y that can be extracted from both *X*_1_ or *X*_2_, Unq() indicates the information that is uniquely disclosed by only *X*_1_ or *X*_2_, and Syn() indicates the information about *Y* that is disclosed by the joint states of *X*_1_ and *X*_2_ together and no simpler combination of elements. In addition to the mutual information between the joint state of *X*_1_ and *X*_2_ and *Y*, the bivariate mutual information can be similarly decomposed:

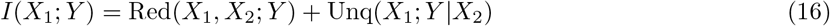

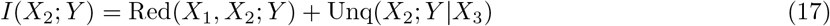

The combination of Eqs. 15, 16, and 17 creates a linear system of equations with three equations, three known values, and four unknown values. If it is possible to determine one of these values (the redundant information, unique information, or synergistic information), the rest can be solved with simple algebra. Unfortunately, classical Shannon information theory doesn’t provide a natural measure of any of them (note that *I*(*X*_1_; *X*_2_) ≠ Red(*X*_1_, *X*_2_; *Y*) and that *I*(*X*_1_; *Y*) ≠ Unq(*X*_1_; *Y*|*X*_2_).

A peculiarity of the PID framework is that, while it provides an elegant structure with which to understand the decomposition of information, it does not specify how exactly to calculate any of the desired values. It is standard to begin with a redundancy function *Red()*, from which the remaining PI-atoms can be recursively calculated, although the question of exactly *what* redundancy function to use is a matter of great contention.

#### 4.9.1 Redundancy / Synergy Bias

In addition to the notion of synergy-as-information-modification, the partial information decomposition also provides insights into what joint information is redundantly shared between parents [40]. It is known that different systems can be variously dominated by redundant or synergistic information dynamics [58, 94], although this is a very new area of study. Previous work has suggested that different behavioral states may be associated with different distributions of redundant and synergistic information processing [59], and we hypothesized that the fixation/cue states would have a different ratio of synergistic and redundant information dynamics. The idea of an explicitly normalized redundancy/synergy ratio has been proposed in [94], who found that topologically similar systems can nevertheless have strongly different distributions of partial information over the PI lattice. The calculate the redundancy/synergy ratio (*RSR*), we first normalize the relevant PI-atoms by the overall joint mutual information, which gives us the proportion of the total information that is purely synergistic and the proportion of the total information that is purely redundant:

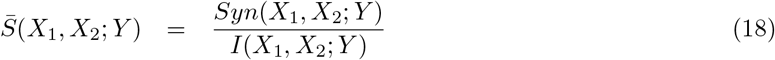

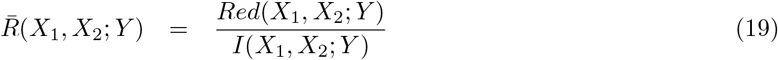

The ratio is then:

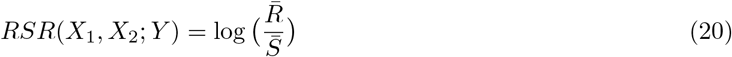

Following the convention established in [58, 94], for the log-redundancy/synergy ratio, a value greater than zero indicates a redundancy-dominated dynamic, while a value less than zero indicates a synergy dominated dynamic.

### 4.10 Data & Code Availability Statement

All scripts used to produce the results described here are available as Supplementary Material, as will all data necessary to replicate the figures and analysis.

